# Comprehensive characterization of the antibody responses to SARS-CoV-2 Spike protein after infection and/or vaccination

**DOI:** 10.1101/2021.10.05.463210

**Authors:** Meghan E. Garrett, Jared G. Galloway, Caitlin Wolf, Jennifer K. Logue, Nicholas Franko, Helen Y. Chu, Frederick A. Matsen, Julie Overbaugh

## Abstract

**Background:** Control of the COVID-19 pandemic will rely on SARS-CoV-2 vaccine-elicited antibodies to protect against emerging and future variants; an understanding of the unique features of the humoral responses to infection and vaccination, including different vaccine platforms, is needed to achieve this goal.

**Methods:** The epitopes and pathways of escape for Spike-specific antibodies in individuals with diverse infection and vaccination history were profiled using Phage-DMS. Principal component analysis was performed to identify regions of antibody binding along the Spike protein that differentiate the samples from one another. Within these epitope regions we determined potential escape mutations by comparing antibody binding of peptides containing wildtype residues versus peptides containing a mutant residue.

**Results:** Individuals with mild infection had antibodies that bound to epitopes in the S2 subunit within the fusion peptide and heptad-repeat regions, whereas vaccinated individuals had antibodies that additionally bound to epitopes in the N- and C-terminal domains of the S1 subunit, a pattern that was also observed in individuals with severe disease due to infection. Epitope binding appeared to change over time after vaccination, but other covariates such as mRNA vaccine dose, mRNA vaccine type, and age did not affect antibody binding to these epitopes. Vaccination induced a relatively uniform escape profile across individuals for some epitopes, whereas there was much more variation in escape pathways in in mildly infected individuals. In the case of antibodies targeting the fusion peptide region, which was a common response to both infection and vaccination, the escape profile after infection was not altered by subsequent vaccination.

**Conclusions:** The finding that SARS-CoV-2 mRNA vaccination resulted in binding to additional epitopes beyond what was seen after infection suggests protection could vary depending on the route of exposure to Spike antigen. The relatively conserved escape pathways to vaccine-induced antibodies relative to infection-induced antibodies suggests that if escape variants emerge, they may be readily selected for across vaccinated individuals. Given that the majority of people will be first exposed to Spike via vaccination and not infection, this work has implications for predicting the selection of immune escape variants at a population level.

**Funding:** This work was supported by NIH grants AI138709 (PI Overbaugh) and AI146028 (PI Matsen). Julie Overbaugh received support as the Endowed Chair for Graduate Education (FHCRC). The research of Frederick Matsen was supported in part by a Faculty Scholar grant from the Howard Hughes Medical Institute and the Simons Foundation. Scientific Computing Infrastructure at Fred Hutch was funded by ORIP grant S10OD028685.

## INTRODUCTION

The future of the COVID-19 pandemic will be determined in large part by the ability of vaccine-elicited immunity to protect against current and future variants of the SARS-CoV-2 virus. Several vaccines have now been approved for use in multiple countries, including two that are based on mRNA technology: BNT162b2 (Pfizer/BioNTech) and mRNA-1273 (Moderna). In the United States, over half of adults are now vaccinated against SARS-CoV-2, the majority of whom have received one of these mRNA vaccines. While these vaccines have been shown to effectively guard against infection, severe disease, and death related to SARS-CoV-2^1-7^, less is known about how effective they will be against emerging and future variants. Current surges in the Delta variant coupled with reports of reduced potency of vaccine elicited antibodies against this variant highlight this concerning ongoing dynamic^8,9^. Evidence from related endemic coronaviruses indicates that evolution in the Spike protein results in escape from neutralizing antibodies elicited by prior infection^10^, potentially contributing to why endemic coronaviruses can reinfect the same host^11-13^. Without immunity that is robust in the face of antigenic drift, continual updates of the vaccine to combat new SARS-CoV-2 variants will likely be necessary to provide optimal protection against symptomatic infection.

Prior infection with SARS-CoV-2 also provides some immunity against subsequent re-infection, and several studies have characterized the epitopes targeted by convalescent sera^14-18^. It is currently unknown whether SARS-CoV-2 infection and vaccination result in antibodies that bind to similar epitopes, an important point to consider given that most people have acquired antibodies through immunization and not infection. The Spike protein encoded by the mRNA in both SARS-CoV-2 vaccines is stabilized in the prefusion conformation by addition of two proline substitutions^19^. This change in sequence and fixed conformation of the Spike protein could result in altered antibody targeting when compared to antibodies elicited during infection, where Spike undergoes several conformational changes. It is also possible that differences in antibody specificity could be due to the amount of antigen or type of immune response stimulated in the context of infection versus vaccination. We know that vaccines drive higher neutralization titers and more Spike binding IgG antibodies than infection^20-22^, indicating some differences in the B cell response compared to infection. A recent study showed that antibodies against the receptor binding domain (RBD) of Spike differ between infected and vaccinated individuals; they are generally less sensitive to mutation and bind more broadly across the domain in the context of vaccination as compared to infection^23^.

Although the majority of the serum binding response in SARS-CoV-2 infected and vaccinated people is directed towards regions of the protein outside of the RBD epitopes^15,23-25^, few studies have examined the prevalence and escape pathways of these antibodies, especially in the setting of vaccination. Antibodies to linear epitopes in the S2 domain of Spike overlapping the fusion peptide (FP), and in the stem helix region just upstream of heptad repeat 2 (SH-H) region are found in serum from COVID-19 patients, and some studies suggest these antibodies may be neutralizing^26,27^. These non-RBD responses may also be important contributors to non-neutralizing antibody activities, which have been associated with protection and therapeutic benefit in experimental SARS-CoV-2 models and with vaccine protection^28-31^. Importantly, these epitopes lie in more conserved regions of Spike than RBD where functional constraints on variation may counter the selective pressure for viral escape.

To compare antibody immunity elicited by SARS-CoV-2 infection and vaccination, we used a high-resolution Spike-specific deep mutational scanning phage display library to profile the epitopes and sites of escape for serum antibodies from people who had been infected, vaccinated, or a combination of both. This approach, called Phage-DMS, identified four non-RBD antibody binding epitopes across all samples: the FP and SH-H region in the S2 subunit, and the N-terminal and C-terminal domains (NTD and CTD, respectively) in the S1 subunit of Spike. Antibodies to NTD and CTD were uniquely present in the setting of mRNA vaccination or severe infection, but mostly absent in mild COVID-19 cases. In vaccinated individuals, the magnitude of the response varied over time both to the CTD and SH-H epitopes. Other covariates, such as age, dose, and vaccine type had no significant differences in the binding profiles observed. Of particular relevance to protection against emerging variants, infection and vaccination appear to shape the pathways of escape differently in different epitopes. In the FP epitope, which is a dominant response after infection, the escape pathway was maintained after subsequent vaccination; in the SH-H epitope, infection resulted in antibodies with diverse pathways of escape, whereas vaccination induced a highly uniform escape profile across individuals. Overall, these findings indicate that vaccination induced a broader antibody response across the Spike protein but induced a singular antibody response at the SH-H epitope, which could favor variants that emerge with these mutations.

## RESULTS

### Samples from individuals with varying SARS-CoV-2 infection and mRNA vaccination histories profiled using high resolution Spike Phage-DMS library

We collected serum samples from two cohorts, termed the Moderna Trial Cohort and the Hospitalized or Ambulatory Adults with Respiratory Viral Infections (HAARVI) Cohort^24,32^. The Moderna Trial Cohort were participants in a Phase 1 trial and consisted of 49 individuals, 34 who received the 100 μg dose of mRNA-1273 (Moderna) and 15 who received the 250 μg dose. Serum samples were taken at days 36 and 119 post first dose (7 and 90 days post second dose, respectively^32^. Serum samples were taken at days 36 and 119 post first dose (7 and 90 days post second dose, respectively)^32^. The HAARVI Cohort included 64 individuals, 44 who had confirmed SARS-CoV-2 infection and 20 who had no reported infection; among this group, 44 were also vaccinated. Those with infection history were stratified by severity based on hospitalization status (39 non-hospitalized/mild vs. 5 hospitalized/severe) and serum was sampled at timepoints ranging from 8 to 309 days post symptom onset. Of these 44 individuals, 24 were also sampled after vaccination with two doses of either mRNA-1273 (Moderna, n=8) or BNT162b2 (Pfizer/BioNTech, n=15), with 23 from the non-hospitalized group and 1 from the hospitalized group. All 20 SARS-CoV-2 naïve individuals were sampled post-vaccination, with 18 having an additional sample taken pre-vaccination (0 to 98 days). Post-vaccination timepoints for all naïve and convalescent individuals ranged from 23 to 65 days after the first dose (5 to 42 days after the second dose, respectively). Figure 1 provides an illustration of the two cohorts and their respective samples’ infection and vaccination statuses. Additional details are available in Supplementary Table 1.

**FIGURE 1:**
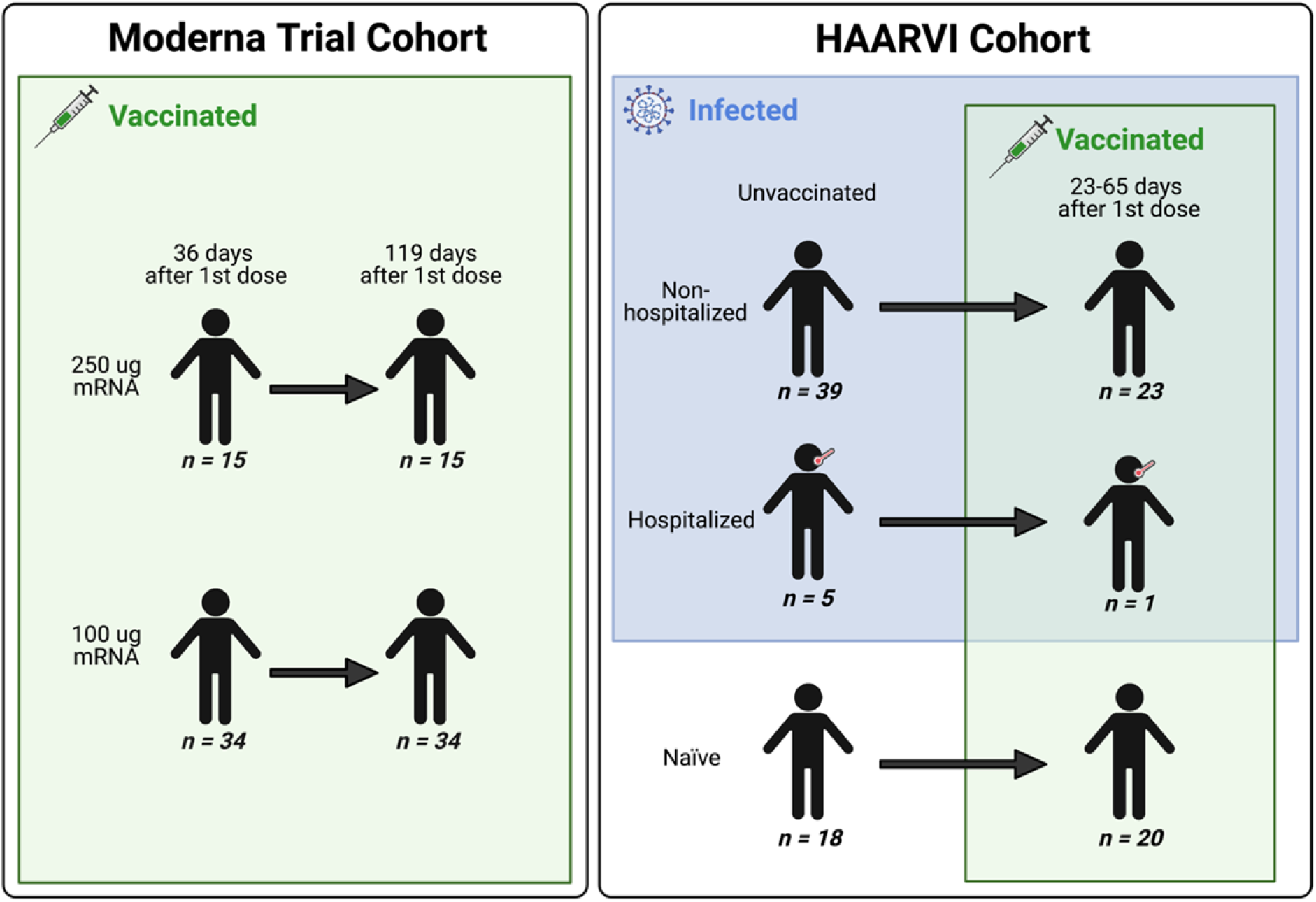
A schematic of sample cohorts. Characteristics of individual participants sampled as part of the Moderna Trial Cohort (left) or the HAARVI Cohort (right). Sample sizes of unique individuals in each group are designated below each figure.

We used a previously described Spike Phage-DMS library to profile the epitopes bound by serum antibodies in the samples described above^24^. This library consists of peptides displayed on the surface of T7 bacteriophage that are 31 amino acids long, tiling across the length of Spike in one amino acid increments. Peptides in the library correspond to the wild-type Wuhan Hu-1 Spike sequence as well as sequences that contain every possible single amino acid mutation at the central position of the peptide. Serum samples were screened with this library by performing immunoprecipitation (IP) followed by sequencing of the pool of phage enriched by the serum antibodies as previously described^24,33,34^.

### Serum antibodies bind to distinct epitopes in infected and vaccinated individuals

We first examined the wild-type peptides in the Spike Phage-DMS library that were enriched by each serum sample to determine the epitopes bound by antibodies in each sample from these cohorts (Figure 2A). The major targeted epitopes across all the cohorts were in the NTD, CTD, FP, and SH-H regions. Serum from non-vaccinated infected individuals who were not hospitalized mostly bound to immunodominant epitopes in the FP and SH-H, both of which are epitopes previously identified in infected individuals using Phage-DMS^24^. Samples from hospitalized/severe COVID-19 cases and vaccinated individuals also bound to the FP and SH-H regions, but additionally bound to epitopes within the NTD and CTD regions. In naïve serum samples there were antibodies that occasionally bound to the FP and SH-H peptides. These findings likely reflect that some individuals have preexisting cross-reactive antibodies that bind to these conserved regions between SARS-CoV-2 and endemic coronaviruses, as suggested by previous studies^16,18^.

**FIGURE 2:**
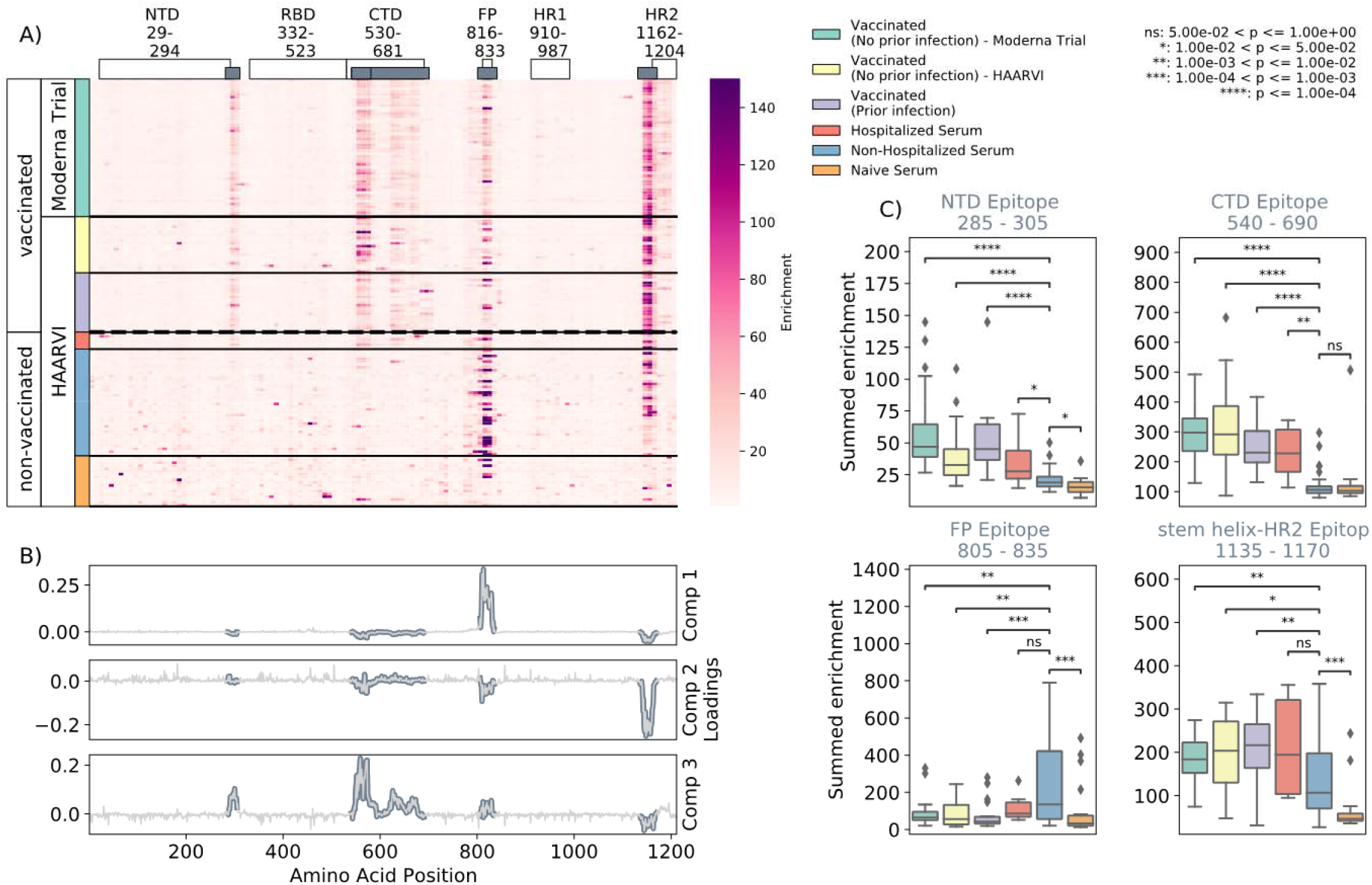
Enrichment of wild-type peptides by serum antibodies. (A) Heatmap with a sample in each row and groups of samples colored on the left. Columns represent peptide locations, with each square on the heatmap indicating the summed enrichment value within a 10-peptide interval. Darker purple indicates higher enrichment values, and values above 150 were capped. Transparent boxes above the heatmap annotate the Spike protein domains, while the smaller grey boxes indicate *epitope binding regions* defined in this analysis (B) The loading vectors from the PCA analysis with the four epitope sites highlighted; enrichments in each of these regions are summed together for subsequent analysis. (C) Box plots describing the distribution of summed wild-type enrichment values for each sample within each of the four epitope sites, each named according to its associated protein domain. Color indicates the sample group. The bars between boxplots give statistical significance (p-value) tests using a Mann-Whitney-Wilcoxon test. All sample group comparisons with the non-hospitalized infected group were performed, and only significant values are shown.

A Principal Component Analysis (PCA) was used to further investigate differences between the infected and/or vaccinated groups. This analysis indicated that binding to epitopes in the NTD, CTD, FP, and SH-H regions were driving differences between samples (Figure 2B). To quantify differences in antibody binding between groups, for each sample we summed together the enrichment values within each identified epitope region and performed pairwise comparisons between non-hospitalized infected people and all other groups (Figure 2C). Most strikingly, we found non-trivial group differences in the magnitude of humoral responses to these major epitopes on the Spike protein. Specifically, antibodies from both hospitalized infected and vaccinated individuals had significantly higher binding to the NTD, CTD, and SH-H regions compared to non-hospitalized infected individuals. However, antibodies from non-hospitalized infected individuals displayed significantly higher binding to the FP epitope than samples from hospitalized or vaccinated individuals. There was no significant difference in any epitope binding in these four regions between vaccinated samples with and without prior infection (p>0.05, Mann-Whitney-Wilcoxon [M.W.W.]).

### Effect of age, dose, vaccine type, and timepoint on epitope binding

In order to determine if there were covariates that contributed to differences in antibody binding, we examined the effect of participant age, vaccine dose and type, and timepoint post infection or vaccination on binding to the four epitopes identified above (Figure 3). For samples in the Moderna Trial Cohort, there was significantly decreased binding to the CTD epitope and SH-H epitope (p=0.008, p=0.011, Wilcoxon rank-sum test with Bonferroni correction) at the later timepoint post first dose (day 119) compared to the earlier timepoint (day 36) (Figure 3A). To examine the effect of dosage, we compared 100 ug and 250 ug mRNA-1273 groups for those between the age of 18 to 55, as that was the only age group included for the 250 ug dose (Figure 3B). There was no significant difference by vaccine dosage for any of the four epitope regions (NTD, CTD, FP, or SH-H). Participant age was also examined as a variable; there appeared to be a difference in epitope binding in the SH-H region, but this did not survive multiple testing correction (Figure 3C).

**FIGURE 3:**
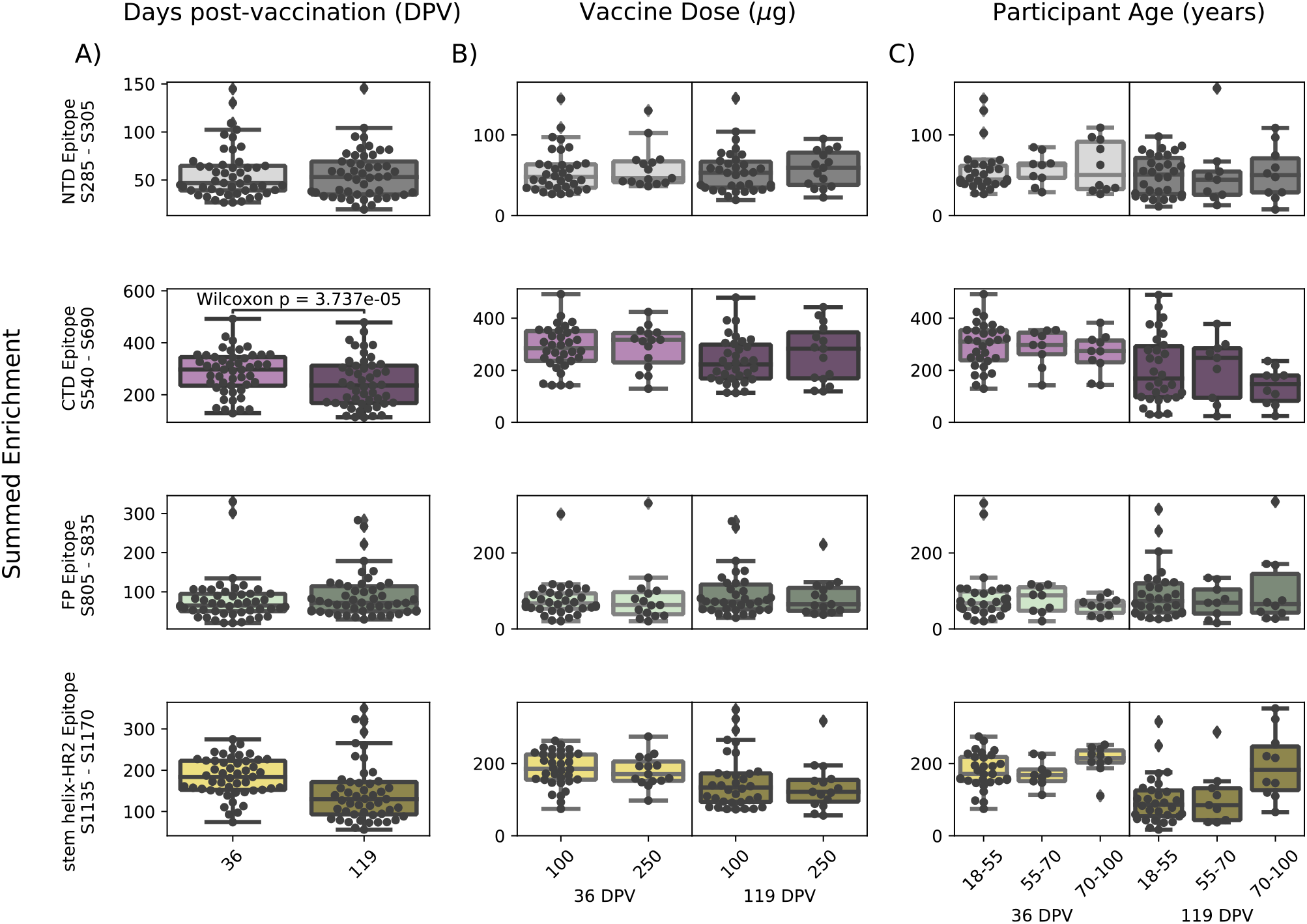
Comparison of epitope binding for NIH Moderna Trial subgroups. Boxplots of summed wild-type enrichment within epitope binding regions for samples grouped by (A) timepoint post vaccination, (B) vaccine dose, or (C) participant age. Samples were taken at either at 36 (n=64) or 119 (n=64) days post vaccination. (B) and (C) are additionally separated by timepoint post vaccination. Results of a Wilcoxon rank-sum test between the groups appears only where p < 0.05 after Bonferroni multiple testing correction (36 group comparisons). Figures containing all p-values for both replicate batches are available at https://github.com/matsengrp/phage-dms-vacc-analysis.

In infected individuals, the effect of time post symptom onset on epitope binding was examined using non-hospitalized infected individuals in the HAARVI Cohort, who were sampled between 26 and 309 days post symptom onset (Supplemental Figure 2A). Samples were binned into three groups: 0-60, 60-180, and 180-360 days post symptom onset. At all times post symptom onset there was no significant difference in binding to the four identified epitopes (p>0.05, M.W.W.). Individuals in the HAARVI Cohort were given either the Moderna mRNA-1273 or Pfizer/BioNTech BNT162b2 mRNA vaccine, and comparison of the epitope binding response between the two vaccine types revealed no significant differences in all epitope regions (Supplemental Figure 2B, p>0.05, M.W.W.).

### Infection and vaccination shape pathways of escape

The Spike Phage-DMS library contains peptides with every possible single amino acid substitution in addition to the wild-type sequence, enabling us to assay the impact of mutations on antibody binding. The effect of site-specific substitutions in critical antibody binding regions not only provides a high-resolution picture of the likely epitope intervals, but also identifies mutations that confer escape within the binding region. The effect of each mutation on serum antibody binding was quantified by calculating its scaled differential selection value, a metric that reports log fold change of mutant binding affinity over wild-type binding affinity at any given site (see Methods)^33^. Site mutations that cause a loss of binding when compared to the wild-type peptide centered at that same site are reported as having negative differential selection values, whereas those that bind better than the wild-type peptide have positive differential selection values. In order for differential selection to be meaningful, however, we must ensure that we do not include weak or sporadic signals that may be due to non-specific binding. Accordingly, we set a threshold of summed wild-type peptide binding in any one region. By doing so, we lose samples in the analysis but can be confident in the results presented by samples passing this curation step (Supplemental Figure 3). For samples that passed this threshold, we compared the effect of prior infection and/or time post vaccination on the pathways of escape in each epitope region as follows. Plots depicting the effect of mutations for all samples are publicly explorable at https://github.com/matsengrp/vacc-dms-view-host-repo.

#### N-Terminal Domain (NTD) and C-terminal Domain (CTD)

We examined the sites of escape within the NTD and CTD epitope regions, focusing on vaccinated individuals from the Moderna Trial Cohort because these epitopes were notable targets of the vaccine response and not commonly found in infected individuals. Vaccination elicited antibodies with a strikingly uniform escape profile in the NTD epitope across samples (Figure 4A), with the majority of samples being sensitive to mutation at sites 291, 294-297, 300-302, and 304, which are in the very C terminal portion of NTD as well as the region between NTD and RBD. The CTD region appeared to consist of multiple epitopes, the dominant being located at the N-terminal region between positions 545 to 580 (termed CTD-N). Antibodies that bound to this dominant CTD epitope had a less uniform escape profile, but sites 561 and 562 were common sites of escape in most samples (Figure 4B). For antibodies to both the NTD and CTD-N epitopes, the pathways of escape tended to drift over time and were different at 119 days post vaccination as compared to 36 days post vaccination.

**FIGURE 4:**
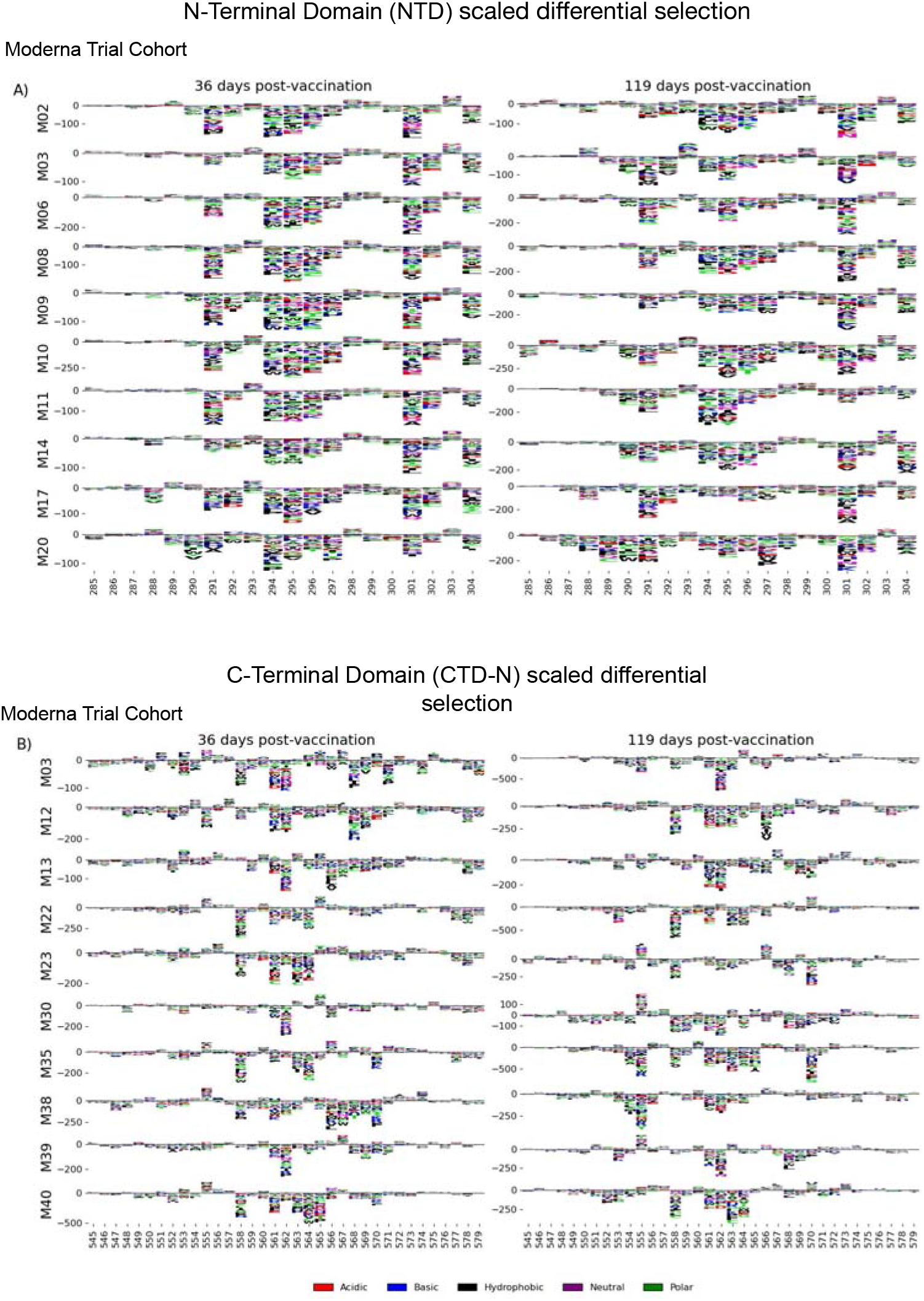
NTD and CTD-N epitope escape profiles. (A and B) Logo plots depicting the effect of mutations on epitope binding in either the NTD (A) or CTD-N (B) epitope for paired samples from the Moderna Trial Cohort. The height of the letters corresponds to the magnitude of the effect of that mutation on epitope binding, i.e. its scaled differential selection value. Letters below zero indicate mutations that cause poorer antibody binding as compared to wild-type peptide, and letters above zero indicate mutations that bind better than the wild-type peptide. Letter colors denote the chemical property of the amino acids. Logo plots on the left and right are paired samples from the same individual, with the participant ID noted on the left.

#### Fusion Peptide (FP)

Antibodies against the FP epitope are strongly stimulated after infection but are less strongly induced after subsequent vaccination (Figure 2). Thus, we investigated whether the pathways of escape for serum antibodies also changed after vaccination within samples from previously infected individuals in the HAARVI Cohort. The escape profiles of antibodies in paired samples that strongly bound to the FP epitope both after infection and after subsequent vaccination are shown as a logo plot (Figure 5A). The major sites of escape within the FP epitope for these samples were sites 819, 820, 822, and 823, and these sites of escape did not appear to change after vaccination although we noted there was more variability in the escape profiles after vaccination.

**FIGURE 5:**
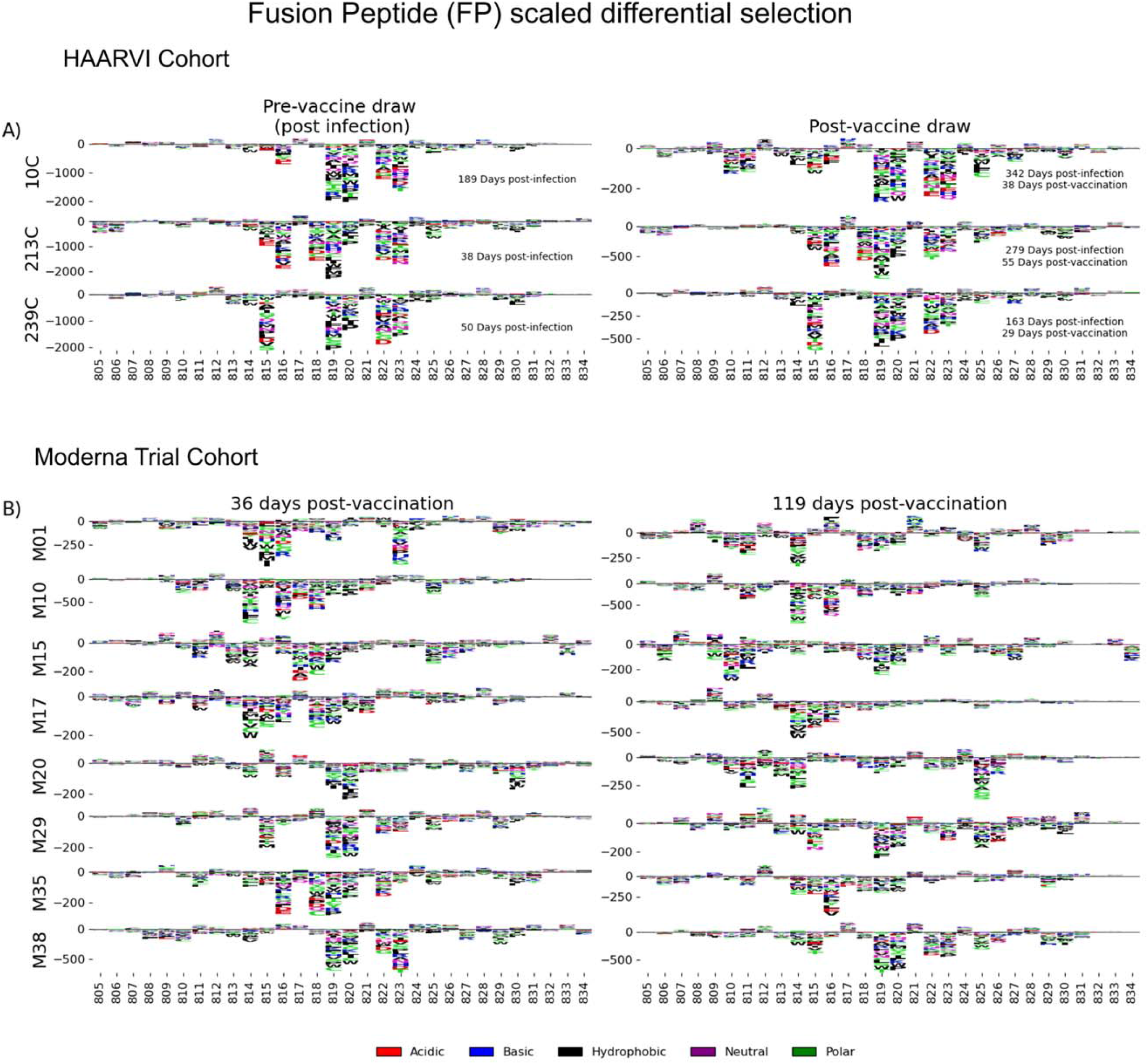
FP epitope escape profiles. (A and B) Logo plots depicting the effect of mutations on epitope binding within the FP epitope region for paired samples from the (A) HAARVI Cohort or (B) Moderna Trial Cohort. Details are as described in Fig 4.

We next examined the pathways of escape for FP binding antibodies in vaccinated individuals from the Moderna Trial Cohort. In people with no prior infection, vaccination induced diverse pathways of escape in the FP region (Figure 5B). For example, for participant M10 escape was focused on sites 814, 816, and 818, whereas for participant M38 escape was focused on 819, 820, and 823. There appeared to be some differences in the escape profile at 119 days as compared to 36 days post vaccination, as exemplified by participants M15, M17, and M20. However, in general many of the major sites of escape were shared at both timepoints within each individual and as a group.

#### Stem Helix-Heptad Repeat 2 (SH-H)

In order to determine the effect of prior infection on the binding profiles of antibodies after vaccination within the SH-H epitope region, we explored the pathways of escape for paired samples from patients with prior infection in the HAARVI Cohort before and after vaccination as was done for the FP region. Samples from previously infected individuals with no vaccination history had diverse pathways of escape within the SH-H epitope (Figure 6A). For example, site 1149 was only sensitive to mutation for participant 217C, and site 1157 was only sensitive to mutations for participants 120C and 146C. In contrast, the samples from vaccinated individuals, regardless of infection history, tended to have a uniform pathway of escape. The most prominent and consistent sites of escape for vaccinated individuals, both with and without prior infection, were at sites 1148, 1152, 1155 and 1156. Of note, the pre-vaccination sample from an individual with prior infection requiring hospitalization (participant 6C) displayed an escape profile highly similar to those from vaccinated individuals, and this escape profile did not change after vaccination.

**FIGURE 6:**
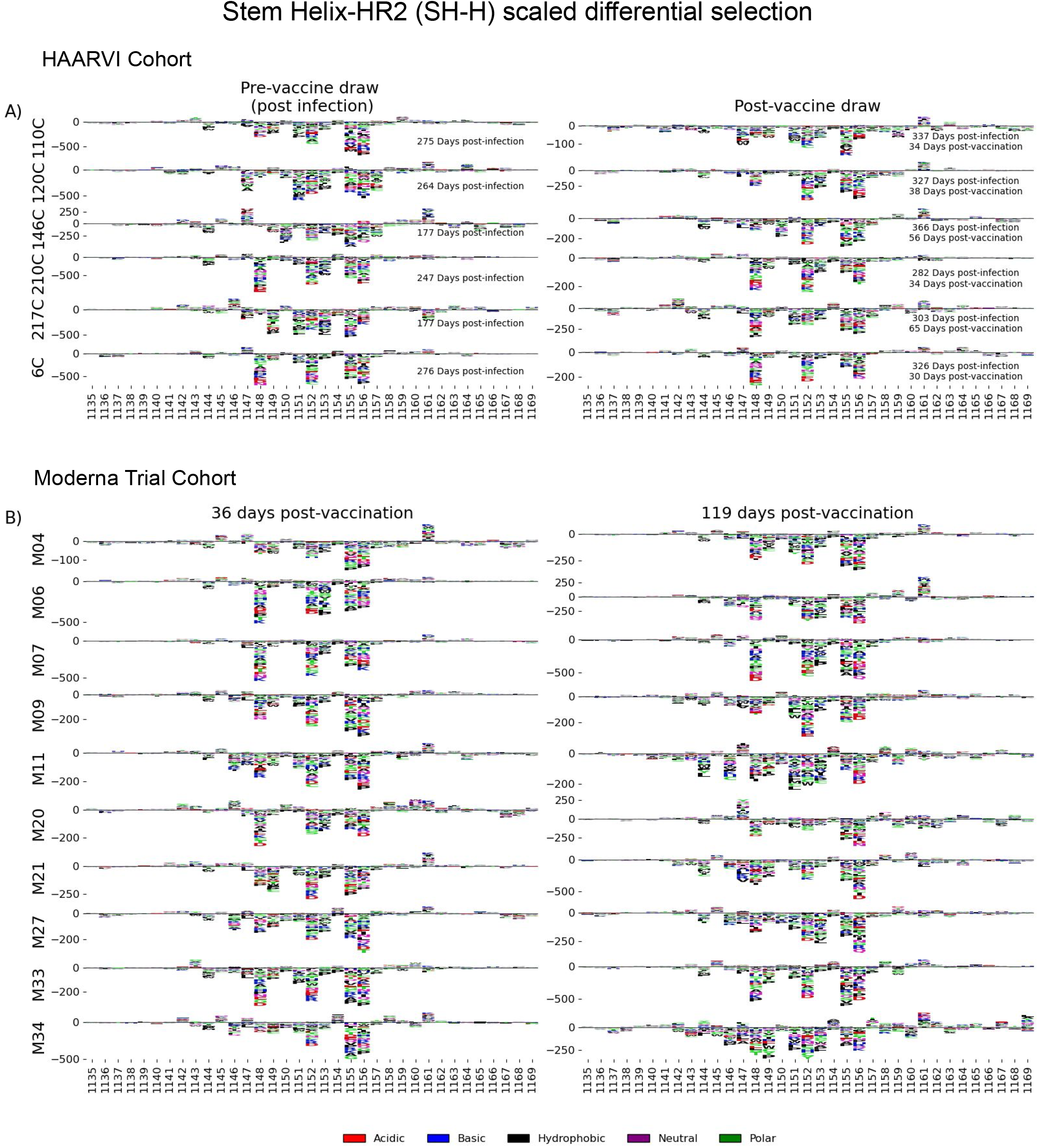
SH-H epitope escape profiles. (A and B) Logo plots depicting the effect of mutations on epitope binding within the SH-H epitope region for paired samples from the (A) HAARVI Cohort or (B) Moderna Trial Cohort. Details are as described in Fig 4.

To see whether the pathways of escape changed over time after vaccination, we visualized the escape mutations within the SH-H epitope for the samples in the Moderna Trial Cohort at 36 and 119 days after the first dose of vaccine (Figure 6B). We saw a highly uniform pattern of escape for most samples at day 36 and 119, again with escape mainly occurring at sites 1148, 1152, 1155 and 1156. For some participants, such as M11, M34, and M35, the escape mutations appeared to drift over time, but the major sites of escape remained the same.

## DISCUSSION

In this study, we comprehensively profiled the antibody response to the SARS-CoV-2 Spike protein, including pathways of escape from sera in individuals with diverse infection and vaccination histories. We identified four major targets of antibody responses outside of the core RBD domains, in the NTD, CTD, FP, and stem helix-HR2 regions. Vaccinated individuals as well as individuals with severe infection requiring hospitalization both had antibodies to these four epitope regions, whereas individuals with mild infection that did not require hospitalization preferentially targeted only FP and SH-H. In previously infected cases, the epitope binding patterns changed over time after vaccination, with decreased binding to both the CTD and SH-H epitopes. However, there was not uniform decay across all four epitopes, indicating that waning antibody titers may not occur for all epitopes equally. Other factors such as vaccine dose (100 ug or 250 ug), vaccine type (BNT162b2 or mRNA-1273), and participant age did not significantly affect the specificity of the antibody response.

We explored the pathways of escape for antibodies binding to these key regions in infected and vaccinated people. We defined for the first time the escape pathways for NTD and CTD-N binding antibodies, epitopes that were commonly found in vaccinated individuals but not in infected individuals. In the case of the NTD epitope, which was located at the C-terminal end of the NTD, escape mutations were uniform and consistent amongst vaccinees, while pathways of escape were more diverse for CTD-N antibodies. Individuals with antibodies that strongly bound the FP epitope had focused escape profiles, with the majority of escape occurring at sites 819, 820, 822, and 823, although the sample size of this group is small (N=3). Vaccination did not greatly alter the escape profile in previously infected individuals, nor did vaccination alone induce a strong or uniform response at the FP epitope. In contrast, antibodies that bind in the SH-H epitope region after infection have diverse pathways of escape, while after vaccination they appear to converge on a more uniform pathway of escape that includes mutations at sites 1148, 1152, 1155 and 1156. Interestingly, these are also the sites of contact for a cross-reactive HR2-specific antibody isolated from a mouse sequentially immunized with the MERS and SARS Spike proteins^35^. This hints that a singular antibody clonotype could be elicited when exposed to a stabilized Spike protein, dominating the response in the SH-H region.

We also observed some drift in the pathways of escape within a single person over time after vaccination. This mirrors findings from a recent study that examined sites of escape for RBD-specific antibodies in serum samples from the same Moderna Trial Cohort as used in this study^23^. Together these results suggest that the B cell response after vaccination with Spike mRNA continues to evolve over time. Multiple studies have demonstrated that SARS-CoV-2-specific B cells undergo continued somatic hypermutation in the months after infection, likely due to antigen persistence^36,37^. Spike antigen has been detected in the lymph nodes at least 3 months after vaccination with BNT162b2, and continued maturation of germinal center B cells could be a possible explanation for the changes in epitope binding we observed^38^. Alternatively, turnover of short-term plasma cells and memory B cells could account for loss of antibody binding to certain epitopes.

Our study has important limitations worth noting. Because the Spike Phage-DMS library displays 31 amino acid peptides, we are unable to detect antibodies that bind to conformational epitopes and/or glycosylated epitopes. This is demonstrated by the lack of observable binding to the RBD region, a domain with complex folding and known target of antibodies from infected and vaccinated individuals. However, prior studies of RBD epitopes have already been reported using an overlapping set of samples from the HAARVI Cohort, however, and together these results paint a more complete picture of epitopes across the Spike protein^15,23^. Finally, we only have 5 individuals within the hospitalized group and this small sample size limits our ability to make conclusions about epitope binding in those with severe infection.

Our finding that vaccinated individuals have a broader response across the Spike protein than infected individuals may have important implications for immune durability against future SARS-CoV-2 variants. Evidence suggests that a polyclonal antibody response that is resistant in the face of multiple mutations is necessary for long-lasting immunity against a mutating viral pathogen^39^. Thus, the polyclonal response to vaccination may provide greater protection from infection than the more focused response after infection. However, the number of epitopes targeted provides just one benchmark and the ability to escape at the population level could also be influenced by the diversity of individuals’ antibody responses at each epitope and thus the likelihood that a single escape mutation could be widely selected. At one S2-domain epitope region (SH-H) vaccination induced uniform sites of escape that may be due to a singular type of antibody that would allow escape by the same mutations for all vaccinated people. However, epitopes in the S2 domain tend to be in highly conserved regions with important functions that constrain the virus’ ability to mutate, making escape from these antibodies less likely than for RBD, where escape is already common. Indeed, mutations in the FP and SH-H epitopes are not arising in the global population of SARS-CoV-2^24^, providing some suggestion that these regions may be constrained^40,41^. Overall, further studies of the functional capacity of these vaccine-elicited antibodies targeting epitopes outside of RBD are warranted, to provide a path towards a polyclonal response to epitopes across the full Spike protein. This comprehensive view may further the goal of a more universal coronavirus vaccine that eliminates the need for continual updates of the SARS-CoV-2 vaccine strain due to mutations in variable regions on Spike.

## MATERIALS AND METHODS

### Sample collection

#### Moderna Trial Cohort

We obtained post-vaccination serum samples via the National Institute of Allergy and Infection Disease that were taken as part of a phase I clinical trial testing the safety and efficacy of the Moderna mRNA-1273 vaccine (NCT04283461)^32^. All samples were de-identified and thus all work was approved by the Fred Hutchinson Cancer Research Center Institutional Review Board as nonhuman subjects research. Trial participants were given either 100 ug or 250 ug doses of the mRNA-1273 vaccine, and serum was sampled from all trial participants at 36 days and 119 days post vaccination. See Table 1 for detailed metadata related to each participant and serum sample.

#### HAARVI Cohort

We obtained plasma samples from individuals enrolled in the Hospitalized or Ambulatory Adults with Respiratory Viral Infections (HAARVI) study conducted in Seattle^24^. Individuals were either enrolled upon PCR confirmed diagnosis with SARS-CoV-2 infection or as control subjects prior to receiving vaccination with either BNT162b2 (Pfizer/BioNTech) or mRNA-1273 (Moderna). See Table 1 for detailed metadata related to each participant and plasma sample. For convenience, all plasma and serum samples in this study are referred to as serum. This study was approved by the University of Washington Institutional Review Board.

### Spike Phage-DMS assay

The Spike Phage-DMS library used in this study contained 24,820 designed peptides that tile across the length of the Spike protein. Peptides are each 31 amino acids long and tile by 1 amino acid increments, and correspond to either the wild-type sequence or a sequence containing a single mutation. Serum samples were profiled using the Spike Phage-DMS library as previously described^24^. Following this method, the Spike Phage-DMS library was diluted in Phage Extraction Buffer (20 mM Tris-HCl, pH 8.0, 100 mM NaCl, 6 mM MgSO_4_) to a concentration of 2.964 × 10^9^ plaque forming units/mL, which corresponds to approximately 200,000-fold coverage of each peptide. 10 uL of serum or plasma was added to 1 mL of the diluted library and incubated in a deep 96-well plate overnight at 4ºC on a rotator. 40uL of a 1:1 mixture of Protein A and Protein G Dynabeads (Invitrogen) were added to each well and then incubated at 4ºC for 4 hours on a rotator. Beads bound to the antibody-phage complex were magnetically separated and washed 3x with 400 uL wash buffer (150 mM NaCl, 50 mM Tris-HCl, 0.1% [vol/vol] NP-40, pH 7.5). Beads were resuspended in 40 uL of water and lysed at 95ºC for 10 minutes. The diluted Spike Phage-DMS library was also lysed to capture the starting frequencies of peptides. All samples were run twice, once each with two independently generated Spike Phage-DMS libraries.

DNA from lysed samples were amplified and sequenced as previously described^24^. Two rounds of PCR were performed using Q5 High-Fidelity 2X Master Mix (NEB). For the first round of PCR, 10uL of lysed phage was used as the template in a 25 uL reaction using primers described in^24^. For the second round of PCR, 2 uL of the round 1 PCR product was then used as the template in a 50 uL reaction, with primers that add dual indexing sequences on either side of the insert. PCR products were then cleaned using AMPure XP beads (Beckman Coulter) and eluted in 50 uL water. DNA concentrations were quantified via Quant-iT PicoGreen dsDNA Assay Kit (Invitrogen). Equimolar amounts of DNA from the samples, along with 10X the amount of the input library samples, was pooled, gel purified, and the final library was quantified using the KAPA Library Quantification Kit (Roche). Pools were sequenced on an Illumina HiSeq 2500 machine using the rapid run setting with single end reads.

### Sample curation and replicate structure

All sample IP’s and downstream analysis were run in duplicate across two separate phage display library batches to ensure reproducibility, with the exception of the four acute samples from hospitalized HAARVI participants which were run in singlicate. All results were cross checked with the set of batch replicates to ensure significance fell within one order of magnitude where applicable. For brevity, we present only figures resulting from the single complete set of batch-specific replicates, however, all figures using the second set of library batch replicates are available (see Code and Data Availability). Additionally, some samples were run with “in-line” technical replicates within the same batch. In the case with more than one technical replicate, we selected the sample with the highest reads mapped from each set of batch replicates for our downstream analysis.

### Short read alignment and peptide counts processing

Samples were aliquoted and sequenced targeting 10X coverage of total sample reads to the peptide library reference. We demultiplexed samples using Illumina MiSeq Reporter software. Post sample demultiplexing, we used a *Nextflow* pipeline to process the peptide counts as well as alignment stats for all samples^42^. The tools and parameters describing the workflow are as follows. The index creation and short-read alignment step were done using *Bowtie2*. During alignment we allowed for zero mismatches in the default seed length of each read (20, very sensitive) after trimming 32 bases from the 3’ end of each 125bp read to match the 93 bp peptides in our reference library^43^. *Samtools* was subsequently used to gather sequencing statistics as well as produce the final peptide counts using the *stats* and *idxstats* modules. Finally, the pipeline collected all reference peptide alignment counts and merges them into a single *xarray* dataset coupling sample and peptide metadata with their respective count. Alignment stats for all replicates are seen in Supplementary Figure S6.

Each of the processing steps described here, as well as downstream analysis and plotting, were run using static and freely available Docker containers for reproducibility. We provide an automated workflow and the configuration scripts defining exact parameters. See Code and Data Availability section for more information.

### Epitope binding region identification

Principal Component Analysis (PCA) via Singular Value Decomposition (SVD) was performed on each set of batch replicates using the scikit-learn package^44^. We first subset our dataset to only include wildtype peptide count enrichments from either infected or vaccinated individuals as input. This curation resulted in the matrix, *X* of size *n* × *p* with *n* biologically distinct replicates and *p* enrichment features across the spike protein. All enrichment values were calculated as a fold change in the frequency for any one sample enrichment over the library control enrichment at the same sites. Each feature was mean centered before performing the PCA such that the covariance matrix of *X* is equivalent to *X*^*T*^*X*/(*n* − 1). We can then use the eigendecomposition, *X* = *USV*^*T*^, to describe the data. The principal axes in feature space are then represented by the columns of *V* and represent the direction of maximum variance in the data. Figure S1 shows three facets of this decomposition; Figure S1A: the unit scaled sample “scores” represented by the columns to visualize sample relationship in principal component space; Figure S1B: component loadings (scaled by the square root of the respective eigenvalues in S); and Figure S1C: the first three principal axes/directions in feature space plotted as a function of the WT peptide feature location on the Spike protein. Together, these provide a visualization of key features in the data used in our downstream analysis. We chose our epitope regions as contiguous regions of nonzero value in the loadings in the first three principal axes.

### Identifying high-resolution pathways of escape

In order to ensure reliable measurements of differential selection of single AA variants compared to the ancestral sequence variant, we threw out samples whose respective sum of wild-type enrichment was below a threshold set for each of the defined binding regions (Figure S4). Once curated, we computed the log-fold change in each of the 19 possible variant substitutions at each site. This metric was then scaled by the average of the wild-type sequence enrichment coupled with both the preceding and following wild-type peptide enrichments at any given site. To evaluate escape at each site, we then sum the differential selection metric as described for each variant at a site to examine a more complete picture of the data defining escape patterns in each sample group.

## CODE AND DATA AVAILABILITY

We provide a fully reproducible automated workflow which ingests raw sequencing data and performs all analyses presented in the paper. The workflow defines and runs the processing steps within publicly available and static Docker software containers, including *phippery* and *phip-flow* described in the Methods section. The source code, Nextflow script, software dependencies, and instructions for re-running the analysis can be found at https://github.com/matsengrp/phage-dms-vacc-analysis.

The generalized PhIP-Seq alignment and count generation pipeline script can be found at https://github.com/matsengrp/phip-flow. A template and documentation for the alignment pipeline configuration is available at https://github.com/matsengrp/phip-flow-template. Finally, we provide a python API, *phippery*, to query the resulting dataset post-alignment that can be found at https://github.com/matsengrp/phippery.

All raw sequencing data was submitted to the NCBI SRA under PRJNA765705. Pre-processed enrichment data is available upon request. Additionally, differential selection data and more can be explored interactively using the dms-view toolkit available at https://github.com/matsengrp/vacc-dms-view-host-repo.

For more information regarding code and data availability, please. For original data from the NIH Moderna trial please see Jackson *et al*^32^, and for information on the HAARVI cohort please contact HYC.

## STATISTICAL ANALYSIS

Estimates of significance presented between group continuous distributions of wild-type enrichment were reported using a *Mann-Whitney Wilcoxon* test with the exception of analysis that included only paired longitudinal samples - such as the comparison of 36-and-119 Days post-vaccination - in this case we used a *Wilcoxon signed-rank* test. Bonferroni correction was applied where applicable and adjusted *P* values < 0.05 were presented as significant. All statistical analysis were done using *Python 3*.*6* and plotted using the *statannot* package found here https://github.com/webermarcolivier/statannot. The static Docker container used for all statistical analysis is publicly hosted at https://quay.io/repository/matsengrp/vacc-ms-analysis.

## ACKNOWLEDGEMENTS

We thank Kevin Sung, Thayer Fisher, and Noah Simon for providing advice regarding computational analyses. We thank Cassie Sather and the Genomics core facility for assistance with sequencing. We thank Laura Jackson (Kaiser Permanente), Chris Roberts, Catherine Luke, and Rebecca Lampley [National Institute of Allergy and Infectious Diseases (NIAID), NIH] for assistance with obtaining the mRNA-1273 phase 1 trial vaccine samples. We also thank all research participants and study staff of the Hospitalized or Ambulatory Adults with Respiratory Viral Infections (HAARVI) study. This work was supported by NIH grants AI138709 (PI Overbaugh) and AI146028 (PI Matsen). Julie Overbaugh received support as the Endowed Chair for Graduate Education (FHCRC). The research of Frederick Matsen was supported in part by a Faculty Scholar grant from the Howard Hughes Medical Institute and the Simons Foundation. Scientific Computing Infrastructure at Fred Hutch was funded by ORIP grant S10OD028685.

## CONTRIBUTIONS

J.O. and F.A.M. conceived the project; M.E.G., J.G., F.A.M., and J.O. led the design of the study; H.Y.C. led the HAARVI study, with C.R.W., J.K.L., and N.F. involved in sample collection. M.E.G. performed all experiments, and J.G. performed all computational and data analyses with F.A.M. advising. M.E.G., J.G., F.A.M, and J.O. wrote the paper with input from all authors.

## ETHICS DECLARATIONS

### Competing Interests

H.Y.C. reported consulting with Ellume, Pfizer, The Bill and Melinda Gates Foundation, Glaxo Smith Kline, and Merck. She has received research funding from Gates Ventures, Sanofi Pasteur, and support and reagents from Ellume and Cepheid outside of the submitted work.

## SUPPLEMENTAL FIGURES

**SUPPLEMENTAL FIGURE 1:**
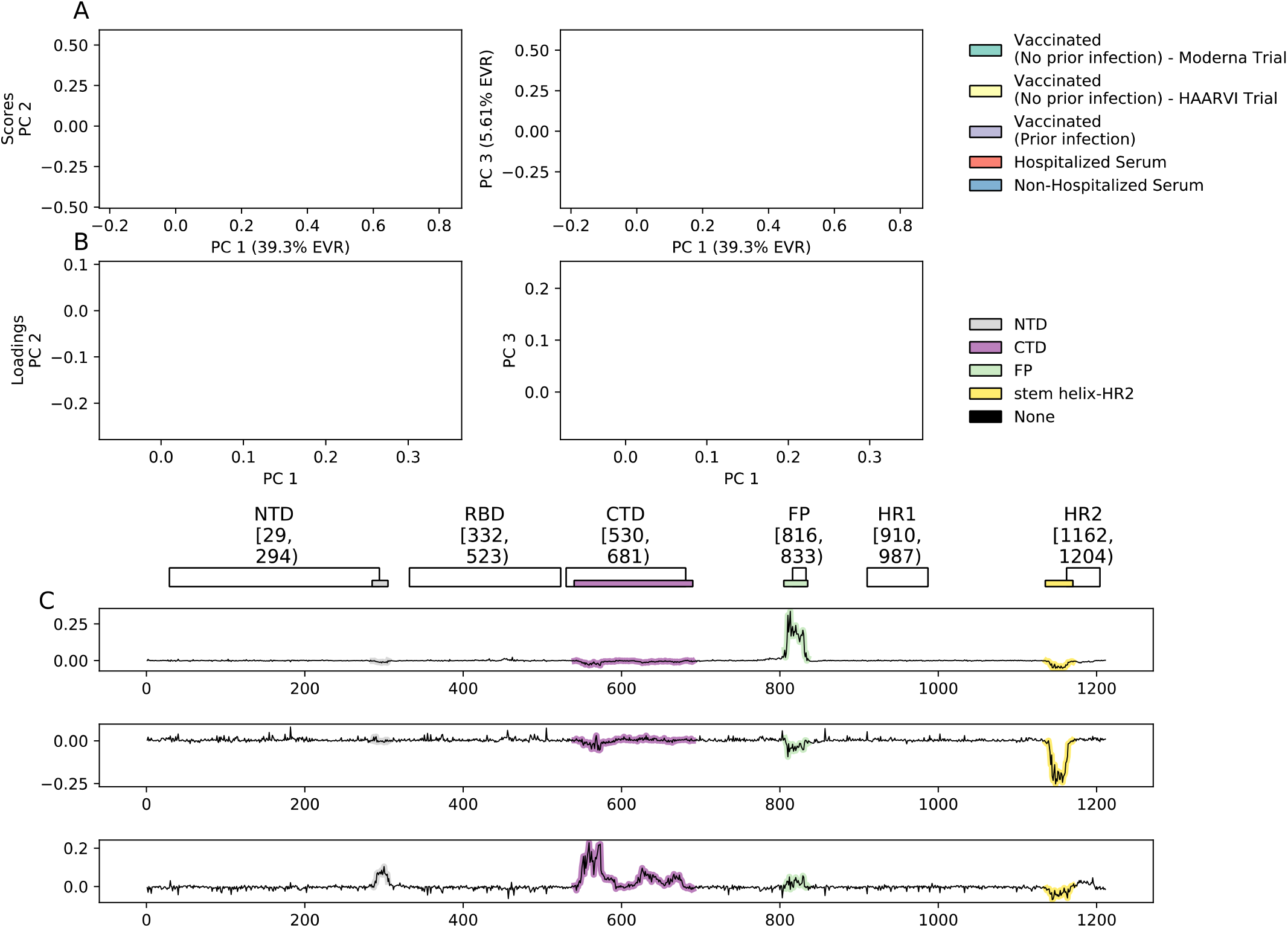
Principal Component Analysis on wild-type enrichment features of all samples. (A) Scatterplot depicting the unit scaled sample “scores” represented by the columns to visualize sample relationship in principal component space. Colors represent the group which each sample belongs to. (B) Vector plots showing the component loadings, scaled by the square root of the respective eigenvalues in the eigen-decomposition. Colors represent the genomic location of each component loading score. (C) Line plots showing the first three principal axes/directions in feature space, plotted as a function of the wild-type peptide feature location on Spike.

**SUPPLEMENTAL FIGURE 2:**
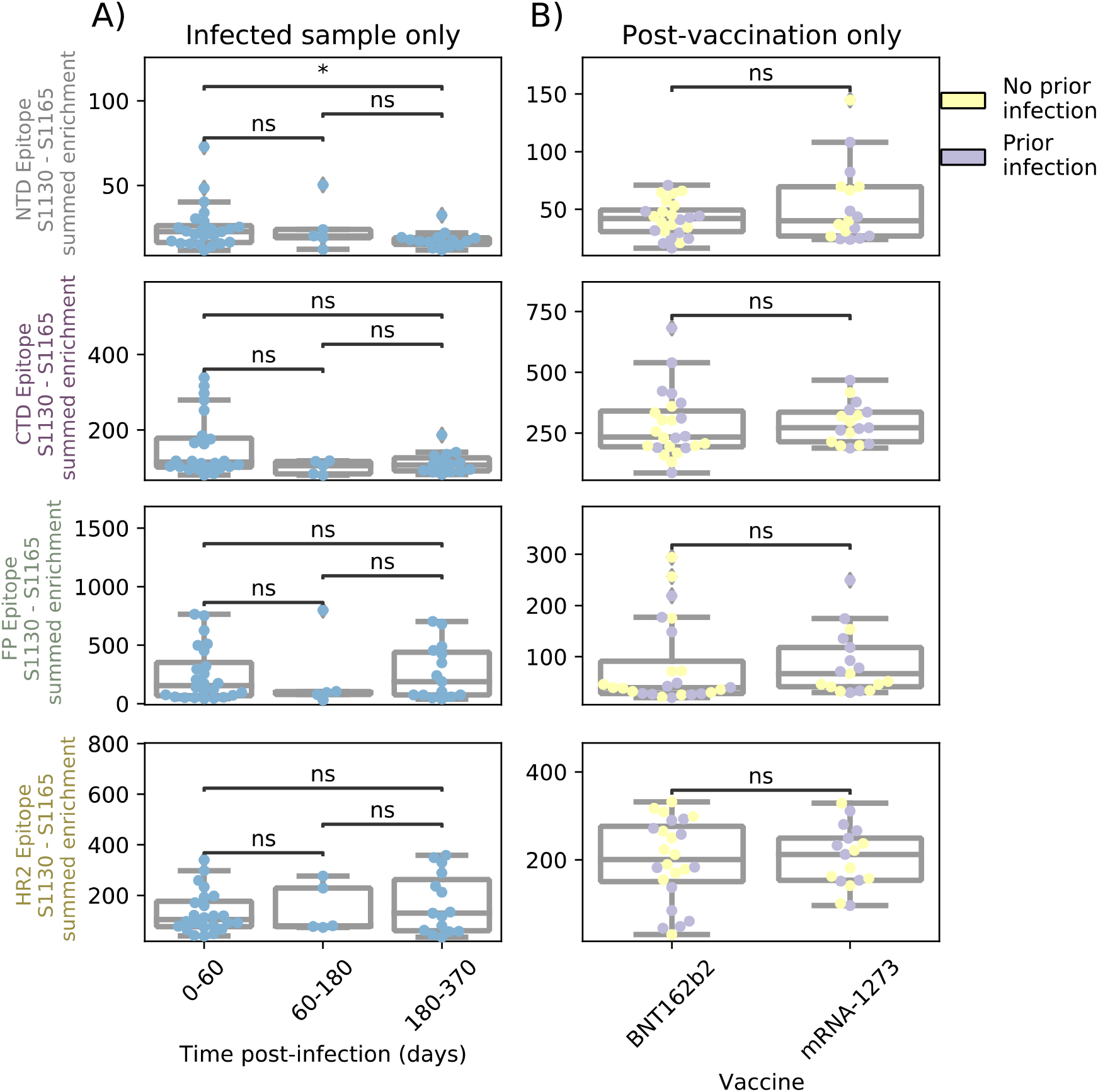
Comparison of epitope binding for HAARVI subgroups. Boxplots of summed wild-type enrichment within epitope binding regions for samples grouped by (A) timepoint post symptom onset or (B) vaccine type (Pfizer/BioNTech BNT162b2 or Moderna mRNA-1273). Results of a Mann-Whitney test between the groups are shown. P-values were adjusted for multiple testing using Bonferroni correction. * indicates p<0.05, ns means “not significant”.

**SUPPLEMENTAL FIGURE 3:**
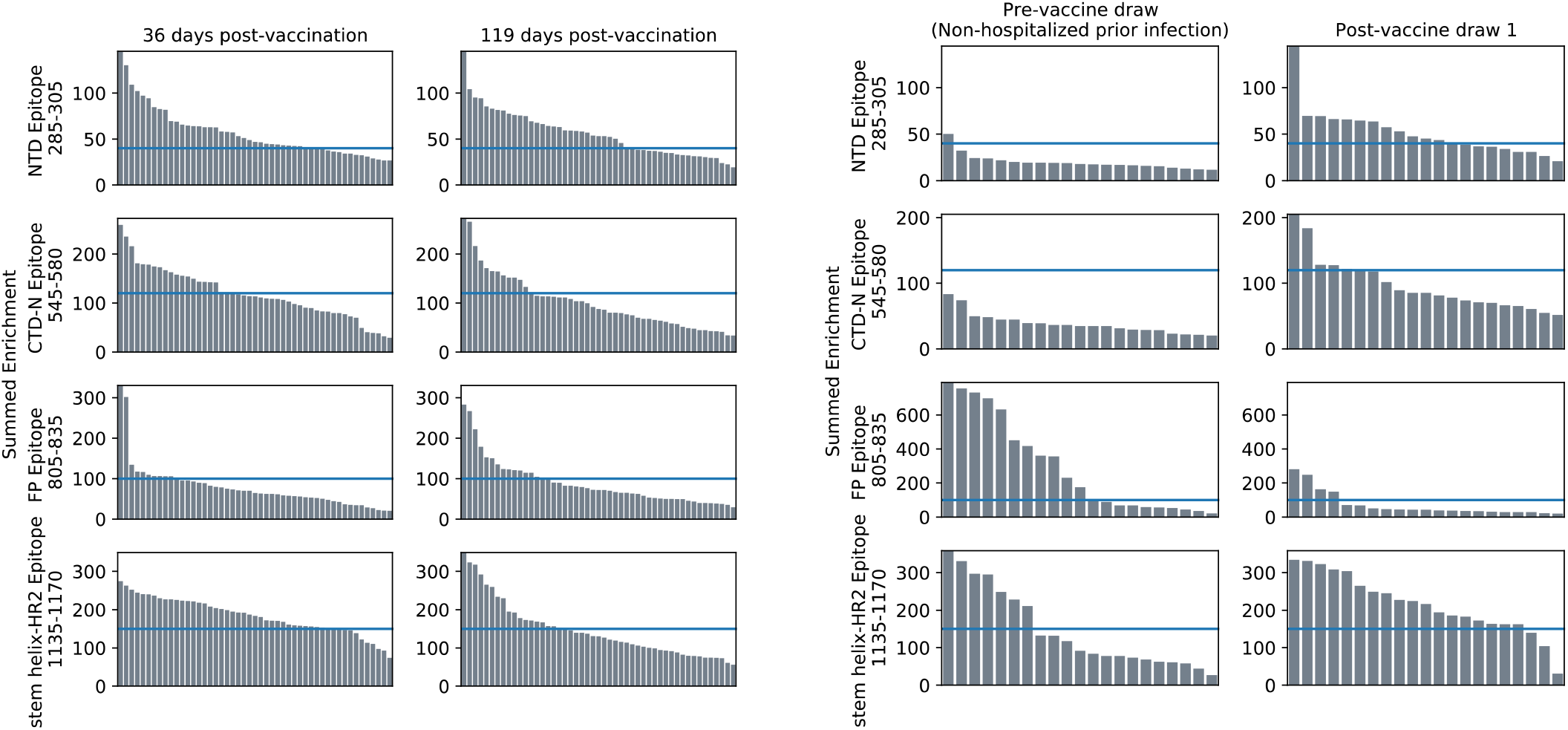
Thresholding of total epitope binding within major epitope regions. Histogram showing the summed enrichment values within each epitope region for every sample in the Moderna Trial Cohort (left two panels) or HAARVI Cohort (right two panels). Blue line delineates the threshold chosen for each epitope region. Samples above the line were included in the escape profile analyses.

## Notes

### Competing Interest Statement

MG and JO are co-inventors on a patent related to Phage-DMS. HYC reported consulting with Ellume, Pfizer, The Bill and Melinda Gates Foundation, Glaxo Smith Kline, and Merck. She has received research funding from Gates Ventures, Sanofi Pasteur, and support and reagents from Ellume and Cepheid outside of the submitted work.

